# PHYLOGENETIC RECLASSIFICATION OF *METARHIZIUM GRANULOMATIS* AND *METARHIZIUM VIRIDE* SPECIES COMPLEX

**DOI:** 10.1101/2025.03.21.644510

**Authors:** Johanna Würf, Volker Schmidt

## Abstract

*Metarhizium* (*M*.*) granulomatis* and *M. viride* have previously been described as pathogens causing hyalohyphomycosis in various species of captive chameleons and bearded dragons (*Pogona vitticeps*). Previous studies yielded five different genotypes of *M. granulomatis* and four different genotypes of *M. viride* based on sequencing of the internal transcribed spacer1-5,8s (ITS1-5,8S) and a fragment of the large subunit of the 28S rDNA (LSU). The aim of this study was to clarify relationships between these genotypes and to obtain a more accurate phylogenetic classification by sequencing of two different loci of the RNA polymerase II second largest subunit, being referred to as RPB1 and RPB2, and the translation elongation factor 1 alpha (TEF). A total of 23 frozen isolates from 21 lizards were available for phylogenetic analysis. 13 isolates belonged to *M. granulomatis*-complex and 10 isolates belonged to *M. viride*-complex. Following the amplification and sequencing of the protein-coding genes, the resulting nucleotide sequences were analyzed, trimmed and assembled. These were further analyzed with regard to differences in single nucleotide polymorphisms (SNPs) and amino acid structure. This was complemented by the construction of phylogenetic trees. The investigation revealed no correlations between the various genotypes and the affected animal species, the isolation site or the clinical/pathological findings. With respect to the present analyses, phylogenetic reclassification is required. Three different genotypes of *M. granulomatis* can be distinguished, which can be phylogenetically addressed as subspecies. Six subspecies can be distinguished regarding *M. viride*.

## 1. Introduction

The importance of fungi within the family of Clavicipitaceae, regarding the genus *Metarhizium* (*M*.) is constantly increasing, especially in reptile medicine (1). As pathogens, these fungi play an important role in poikilothermic animals. The reason for this is, that the immune system of reptiles is heavily dependent on their husbandry conditions, such as the temperature in their enclosure (2). Ongoing, humidity, diet and UV-light conditions have an impact on reptilian health (3). Diseases caused by *Metarhizium* have so far been detected in several species of lizards, turtles, tortoises but also in crocodiles (1). Over the last six decades, various studies have been carried out to gain a better understanding of morphology, pathogenesis and transmission routes (4). In addition, molecular biological studies, such as polymerase chain reaction and DNA sequencing, play a major role in defining and phylogenetically specifying species within this order. (5,6).

*Metarhizium granulomatis* and *M. viride* are known to be a cause of hyalohyphomycosis in captive lizards (Schmidt et al., 2012, 2017b, 2017a, 2018; Klasen et al., 2019). In particular, *M. granulomatis* and *M. viride* have been described as primary pathogens, affecting chameleons and bearded dragons and causing fungal dermatitis, granulomatous glossitis, pharyngitis, as well as disseminated visceral mycosis (Sigler et al., 2010; Schmidt et al., 2012, 2017b). *M. granulomatis* thereby only seems to cause manifest infection in veiled chameleons (*Chameleo calyptratus*). So far there have been studies on medication, but successful treatment of infected lizards has not been achieved yet (7). The route of entry into the enclosure and infection mechanisms still remain unclear and need further investigation. Latest research results suggest that these keratinophilic fungi represent species complexes within which further differentiation may be possible. According to Schmidt et al. (2017b), five different genotypes for *M. granulomatis* (A-E) could be assumed, under which genotype A showed a strong correlation with dermatitis. Genotype B – E therefore could be associated with pharyngitis and glossitis (7). Furthermore, four different genotypes for *M. viride* (A-D) could be assumed (8). However, ribosomal genes of the internal transcribed spacer1-5,8s (ITS1-5,8S) and a fragment of the large subunit of the 28S rDNA (LSU) have been sequenced (Schmidt et al., 2017a,b), which does not appear to be sufficient for phylogenetic studies in the family Clavicipitaceae (5,6).

The objective of this study was to elucidate the relationships among potential subspecies by concentrating on the sequencing of the following protein-coding nucleotides: two different loci of the RNA polymerase II second largest subunit, being referred to as RPB1 and RPB2 and the translation elongation factor 1 alpha (TEF). By evaluation of single nucleotide polymorphisms (SNPs) and translating nucleotides into amino acid pattern, conclusions were drawn to get a better understanding of the phylogeny of *M. granulomatis-* and *M. viride*-complex. The importance of a correct phylogenetic classification thus lies in the potential to discover possible differences in virulence, pathogenesis or preventive measures. It could also provide insight into the host specificity of the fungi.

## 2. Materials and Methods

### 2.1. Fungal isolates

Fungal isolates were collected between 2010 and 2023 from lizards (n = 21) which were presented to the Clinic for Birds and Reptiles, University Leipzig, Germany. The isolates were derived from the following species: veiled chameleons (n = 11), panther chameleons (*Furcifer pardalis*) (n = 4), Parson’s chameleons (*Calumma parsonii*) (n = 3), central bearded dragons (*Pogona vitticeps*) (n = 2), and a carpet chameleon (*Furcifer lateralis*) (n = 1). Furthermore, they were obtained from throat (n = 9), cloaca (n = 7), tongue (n = 4), liver (n = 2), faecal samples (n = 2) or rectum (n = 1). The storage of isolates was in ROTI**®**Store yeast cryotubes (Carl Roth GmbH & Co.KG, Karlsruhe, Germany) at a temperature of -32°C until utilization.

Purification of genomic DNA was done using the DNeasy blood and tissue kit (Qiagen, Hilden, Germany) following the protocols given by the manufacturer. In the following steps species affiliation was proven by ribosomal DNA sequencing (Schmidt et al., 2017a,b).

Regarding *M. granulomatis*, genotype A was obtained from Parson’s chameleons (n = 2) and a panther chameleon, genotype B isolates were obtained from four chameleon species, including one Parson’s chameleon with co-isolation of genotype A. Genotype C and D were obtained only from veiled chameleons (n = 2, each). One of those veiled chameleons revealed genotype B and C. Formerly described genotype E was not included in this study.

With view to *M. viride*, genotype A came from veiled chameleons (n = 2), genotype B from a central bearded dragon and a panther chameleon (one each), genotype C from one veiled chameleon, one panther chameleon and one Parson’s chameleon. Genotype D (n = 1) was obtained from a central bearded dragon. In addition, two isolates with previously unpublished genotyping were obtained from veiled chameleons (n = 2). Until further examination the purified genomic DNA of these isolates was frozen by -32°C in Eppendorf tubes.

### 2.2. PCR and DNA sequencing

The primers employed in this study were selected based on previous phylogenetic attempts to reclassify *Metarhizium* spp. and are listed in Table 1 (Liu et al., 1999; Bischoff et al., 2006; Malkus et al., 2006; Mongkolsamrit et al., 2020).

**Table 1.**
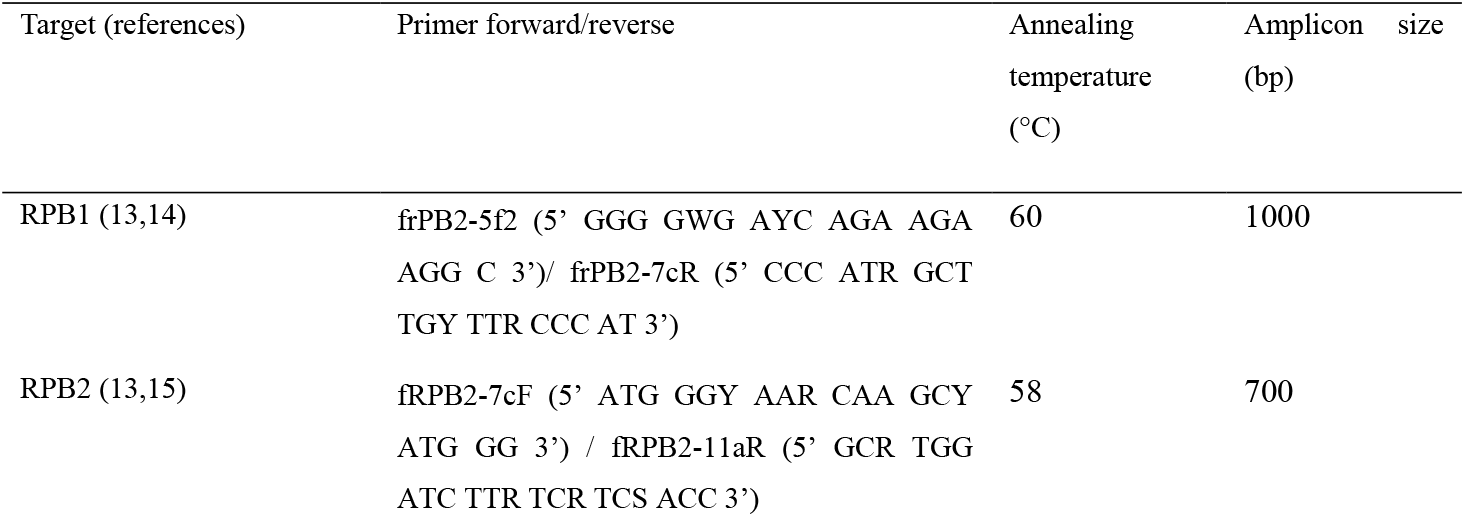
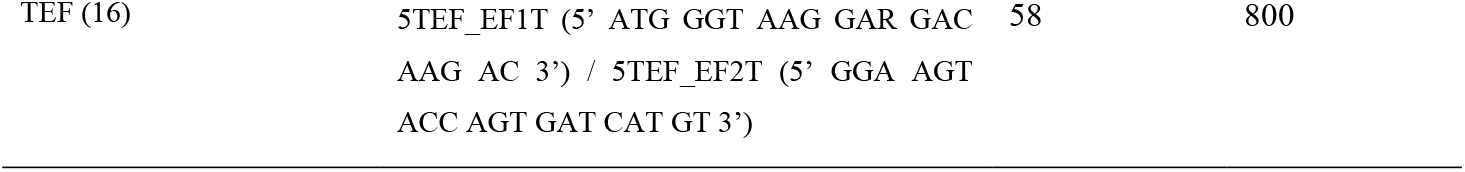
Primers used for amplification and sequencing of two different loci of the RNA polymerase II second largest subunit (RPB1, RPB2) and the translation elongation factor 1 alpha (EF-1alpha) of *Metarhizium granulomatis* and *M. viride*.

All of the samples analyzed according described protocols, focused on partial sequences of three gene loci: RPB1, RPB2 and TEF. (13–16). After samples were amplified, detection was done by gel electrophoresis. PCR products were further processed by Polyethylen Glycol precipitation (17). Sequencing was done with Sanger sequencing by a commercial DNA sequencing service (Faculty of Medicine, Central Functional Area DNA-Technologies / Sequencing, Leipzig, Germany), using 10 μl of sample as well as primers simultaneously as described earlier.

### 2.3. Analyzing data according to phylogeny

The alignment length for each gene locus was 819 bp for RPB1 and 802 bp for RPB2. With respect to TEF it was 739 bp regarding *M. granulomatis* and 819 bp regarding *M. viride*. This length polymorphism was due to various gaps. After alignment, introns were removed from the sequences as the main focus here laid on the coding sequences (CDS). This resulted in a length of 257 bp for the TEF of both *Metarhizium* species. As a next step all sequences were edited in a multilocus dataset with a length of 1878 bp total by using MEGA11 (Molecular Evolutionary Genetics Analysis version 11, Tamura et al., 2021). Comparison with already available data in the GenBank database was being carried out according to the BLAST program (http://www.ncbi.nlm.nih.gov/BLAST/) (19).

Further, the isolates were evaluated in the four following datasets: RPB1, RPB2, TEF and the multilocus dataset. After all isolates were analyzed in regard of single nucleotide polymorphisms (SNPs), they were translated into amino acid pattern to detect any differences in branching that could result therefore. Evolutionary history and potential similarity were derived by Maximum Likelihood method based on Tamura-Nei model (20). Rates among sites were set at Gamma Distributed (G) with a number of 5 discrete Gamma categories. Codon positions included were the first, second, third and noncoding positions. The percentage of the trees in which associated taxa clustered is shown next to the branches (1000 replicates). To create additional outgroup taxa, isolates of the *Metarhizium anisopliae*-group (designated informally as the “PARB” clade) (21) such as *M. anisopliae* [NCBI Acc.-Nr. DQ463996.2 (22)], *M. brunneum* [NCBI Acc.-Nr. EU248854.1 (22)] and *M. robertsii* [NCBI Acc.-Nr. DQ463994 (22)] were added. A member representing the family of *Clavicipitaceae* that was used was *M. acridum* [NCBI Acc.-Nr. MK391183 (23)]. Also, a *Beauveria bassiana* isolate [NCBI Acc.-Nr. MN026878 (Dalla Nora, 2019, unpublished)] was added to act as a root. To underline the results given by the Maximum Likelihood method, pairwise distance was computed by MEGA program using the Maximum Composite Likelihood (MCL) approach.

## 3. Results

### 3.1. RNA polymerase II second largest subunit (RPB1)

Sequencing of the RPB1 showed no differences between the *M. granulomatis* isolates type A, C and D, which were 100% identical to *M. granulomatis* [NCBI Acc.-Nr. KJ398688.1 (6)]. The only difference to *M. granulomatis* type B was one single nucleotide polymorphism. However, translation into amino acid structure showed 100% identity of all *M. granulomatis* isolates as well as to the additional outgroup taxa of *M. anisopliae* [NCBI Acc.-Nr. OK336701.1 (Du, 2021, unpublished)], *M. brunneum* [NCBI Acc.-Nr. EU248934.1 (22)], *M. robertsii* [NCBI Acc-Nr. DQ468368.1 (22)] and *M. acridum* [NCBI Acc.-Nr. KM527862.1 (Zhang, 2014, Unpublished)] regarding the RPB1 (Suppl.1).

Regarding *M. viride*, the isolates of genotype A, genotype D and the two isolates of previously unpublished genotype differed in one single nucleotide each. Translation into amino acids, however, showed that they were 100% identical, with an identity of 100% to M. *viride* [NCBI Acc.-Nr. KJ398717 (6)]. Genotype C can be differentiated from other isolates by three SNPs. This resulted in an amino acid identity of 99.63% to genotype A and D isolates, as well as to the isolates of previously unpublished genotype. Ongoing, two isolates, which were labeled as *M. viride* genotype B, could be distinguished from each other on the hand of four single nucleotide polymorphisms. These differences also translated into amino acid structure, resulting in an identity of 99.63%. One of these isolates [NCBI Acc.-Nr. PV231550] showed an identity of 98.13 – 99.63%, while the other isolate [NCBI Acc.-Nr. PV231551] had an identity of 98.51 – 99.63% compared to the remaining *M. viride* isolates.

*M. granulomatis* [NCBI Acc.-Nr. KJ398688.1 (6)] showed an identity of 99.63% to *M. viride* [NCBI Acc.-Nr. KJ398717 (6)]. Either way, between the different genotypes of *M. viride* and *M. granulomatis* considered in this study, an identity between 98.13 – 99.63% could be observed. The same percentage of identity was shown between *M. viride* and the additional outgroup taxa.

### 3.2. RNA polymerase II second largest subunit (RPB2)

By analyzing the nucleotide structure of RPB2 of *M. granulomatis* isolates, several SNPs could be observed. In total, there were six SNPs regarding genotype B, four regarding genotype C and D and only one SNP regarding genotype A. Neither of these had an impact on amino acid structure, so that all isolates of *M. granulomatis* had an identity of 100%. They also clustered with the additional outgroup taxa of *M. anisopliae* [NCBI Acc.-Nr. OK336701.1 (Du, 2021, unpublished)], *M. brunneum* [NCBI Acc.-Nr. XM_014688283.1 (24)], *M. robertsii* [NCBI Acc.-Nr. XM_007819877.1 (25)] and *M. acridum* [NCBI Acc.-Nr. XM_066120576.1 (26)], likewise with an identity of 100% (Suppl.2).

Regarding *M. viride* isolates, similar correlations regarding RPB2 could be identified. Even though there was a total of nine variable sides on a nucleotide base, no differences in amino acid structure resulted, so that there was an identity of 100% between all genotypes. Ongoing, 100% of identity could be observed between *M. granulomatis* and *M. viride* isolates. In this dataset, a comparison with *M. granulomatis* and *M. viride* isolates from the BLAST database was not possible due to differences in length of the sequenced fragments or because they were not available at all. The *B. bassiana* isolate [NCBI Acc.-Nr. LC812020.1 (Yang et al., 2024, unpublished)] clustered phylogenetically outside with 96.34% identity to the other isolates.

### 3.3. Translation elongation factor 1 alpha (TEF)

Looking at *M. granulomatis* with respect to the nucleotides of the TEF, sixteen variable sites were detected. Only four of these were located on the CDS regions, so that further analyzation focused on them. One of these SNPs concerned isolates belonging to genotype A. This also had an impact on amino acid structure, resulting in genotype A isolates clustering separately from others with an identity of 98.80%, as well as an identity of 100% to *M. granulomatis* [NCBI Acc.-Nr. MH619511.1 (9)]. Ongoing, there were two SNPs in the CDS nucleotide sequence of genotype B isolates, two regarding VS14512 [NCBI Acc.-Nr. PV231585] and VS18409 [NCBI Acc.-Nr. PV231588] and one regarding VS15807 [NCBI Acc.-Nr. PV231586]. These SNPs did not translate into further changes, so that all of the other *M. granulomatis* isolates clustered together with an identity of 100% (Suppl. 3).

The TEF of *M. viride* isolates showed 17 variable sites from which only one was located in the CDS region. This SNP was belonging to the sequence of genotype A isolates. No impact on amino acid structure was observed, so that all isolates clustered together. Maximum likelihood analyses confirmed a 100% identity to *M. viride* [NCBI Acc.-Nr. MH619515.1 (9)].

A total of 78 variable sites were identified between *M. granulomatis* and *M. viride*. Only eight of these single nucleotide polymorphisms were located on the coding regions. Either way, by analyzing these CDS, all isolates apart from *M. granulomatis* genotype A, had an identity of 100%. Isolates of *M. brunneum* [NCBI Acc.-Nr. EU248854.1 (22)], *M. robertsii* [NCBI Acc.-Nr. DQ463994.2 (16)], *M. anisopliae* [NCBI Acc.-Nr. DQ463996.2 (16)]and *M. acridum* [NCBI Acc.-Nr. MK391183.1 (23)] clustered phylogenetically apart as additional outgroup taxa, resulting in an identity of 96.36 – 97.58%.

### 3.4. Multilocus analysis

A total of sixteen variable sites were identified through the evaluation of single nucleotide polymorphisms present within the multilocus dataset of *M. granulomatis*. Genotype A isolates showed a total of eight SNPs, while isolates belonging to genotype C showed five. No differences between genotype C and D could be observed. Furthermore, isolates belonging to genotype B showed eight SNPs. Two of them made it possible to differentiate within this genotype. After creating a phylogenetic tree on nucleotide base, VS15807 clustered separately, furthermore VS18409 and VS14512 clustered together. The SNPs resulting in this branching were located on the sequence of TEF. However, this did not have any impact on amino acid structure, as isolates belonging to genotype B had an identity of 100%. Concerning the amino acids, genotype A showed an identity of 98.69 – 99.02% to other isolates, while genotype C showed an identity of 98.35 – 99.02%. Genotype C and D isolates were 100% identical (Fig.1).

**Fig. 1.**
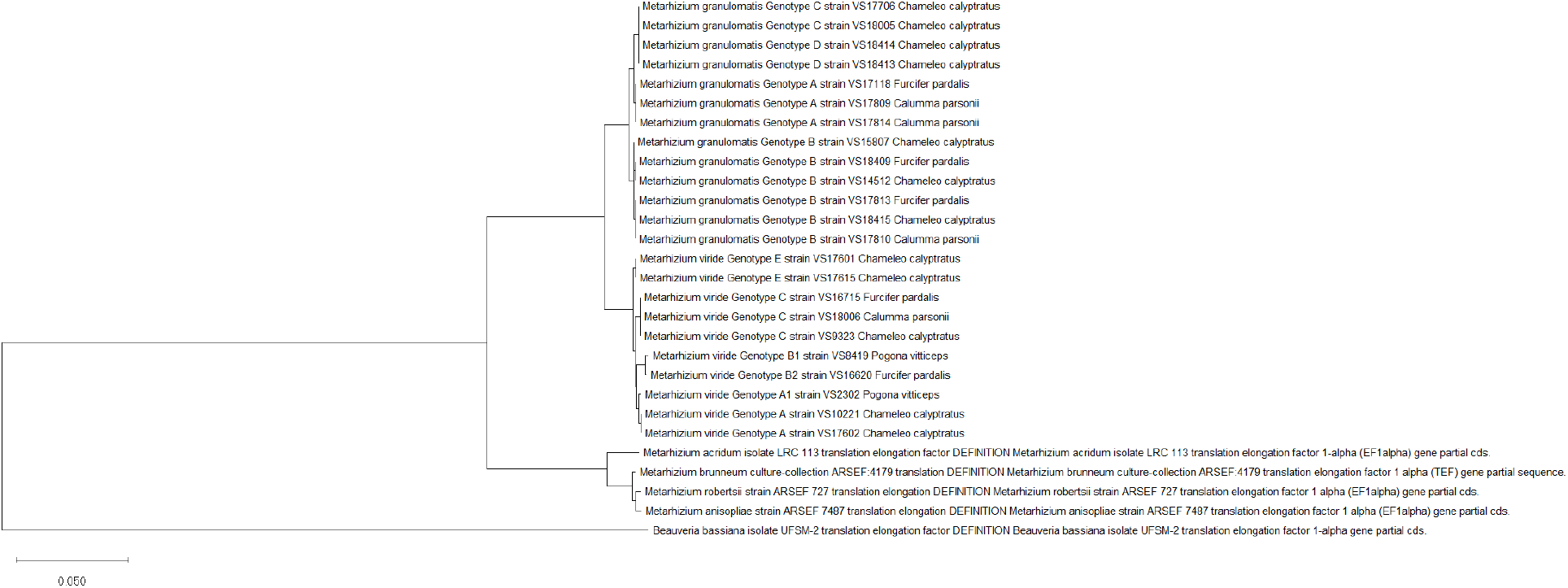
Phylogeny inferred from the analysis of CDS regions of RPB1, RPB2 and TEF from 13 isolates of *Metarhizium* (*M*.) *granulomatis* – complex and 10 isolates of *M. viride* – complex, assembled into a multilocus dateset. Analysis was done based on nucleotide structure. Other isolates of *Claviciptitaceae* (Sordariomycetes: Hypocreales) and *Beaveria bassiana* were added as additional outgroup taxa. Reference sequences with accession numbers were taken from the NCBI GeneBank database (http://www.ncbi.nlm.nih.gov). Phylogeny is based on Maximum likelihood analysis. The tree is drawn to scale, with branch lengths measured in the number of substitutions per site.

With regard to *M. viride*, single nucleotide polymorphisms were numerous and resulted in twenty-five variable sites. As a conclusion, six different branches were built by constructing Maximum Likelihood trees. Genotype A isolates VS17602 and VS10221 differed from VS2302, which was formerly named as genotype D, based on four changes in nucleotide structure. Genotype B isolate VS8419 showed a total of fourteen SNPs, whereas genotype B isolate VS16620 showed twelve. Ongoing, differentiation between the two of them was possible on the hand of four SNPs. Isolates belonging to genotype C showed eight SNPs, likewise did the ones with previously unpublished genotype.

Regarding amino acid sequence, once again it was possible to differentiate within six different branches. Genotype A isolates VS17602 and VS10221 had an identity of 99.51% to the genotype D isolate VS2302, which caused this isolate to cluster separately. Ongoing, this specific genotype showed an identity of 98.18 – 98.85% to remaining isolates. The SNPs of genotype B isolates VS8419 and VS16620 likewise had an impact on amino acid structure, as they had an identity of 99.51%. VS8419 showed an identity of 97.85% to genotype C, as well as 98.18 – 98.35% other isolates, whereas VS16620 showed an identity of 98.02% to genotype C and once again 98.18 – 98.35% to other isolates. The two isolates of previously unpublished genotype had an identity of 99.02% to genotype A, also they showed 98.18 – 99.02% of similarity to other isolates (Fig.2).

**Fig. 2.**
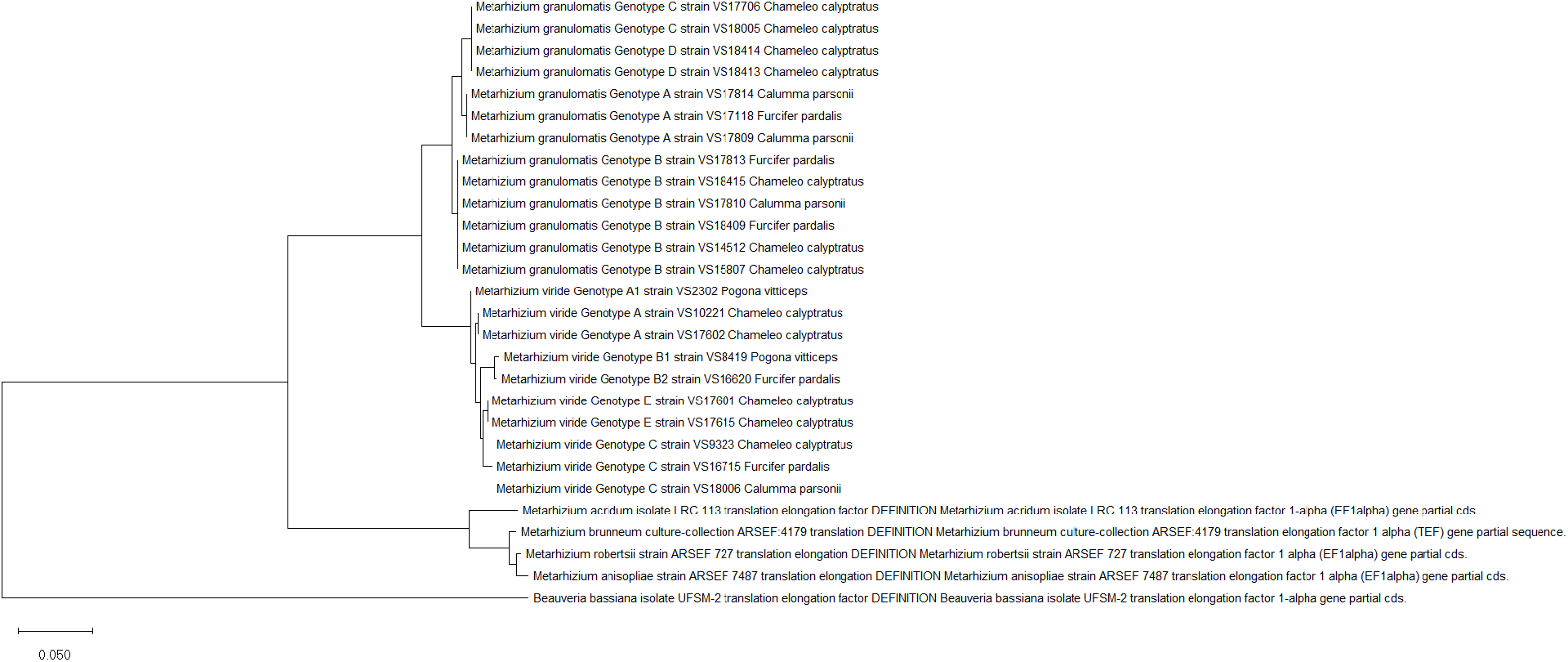
Phylogeny inferred from the analysis of CDS regions of RPB1, RPB2 and TEF from 13 isolates of *Metarhizium* (*M*.) *granulomatis* – complex and 10 isolates of *M. viride* – complex, assembled into a multilocus dateset. Analysis was done based on amino acid structure. Other isolates of *Claviciptitaceae* (Sordariomycetes: Hypocreales) and *Beaveria bassiana* were added as additional outgroup taxa. Reference sequences with accession numbers were taken from the NCBI GeneBank database (http://www.ncbi.nlm.nih.gov). Phylogeny is based on Maximum likelihood analysis. The tree is drawn to scale, with branch lengths measured in the number of substitutions per site.

All *M. granulomatis* isolates could be distinguished from *M. viride* isolates with an identity of 92.53 – 94.46%. Once more, the inclusion of *M. granulomatis* or *M. viride* isolates from the BLAST dataset for comparison was rendered unfeasible, owing to the paucity of extant published data. Maximum likelihood analysis furthermore confirmed that the earlier described PARB clade is distinct from *M. granulomatis* and *M. viride* isolates, with *M. granulomatis* isolates having an identity of 73.12 – 73.97%. *M. viride* isolates showed an identity of 71.35 – 73.93% to the outgroup taxa. *B. bassiana* [NCBI Acc.-Nr. MN026878.1 (Dalla Nora, 2019, unpublished)] once again functioned as an outgroup.

### 3.5. Isolates and associated findings

Further evaluation concerning the source of isolation revealed that isolates belonging to *M. granulomatis* genotypes A and B could all be obtained from cloacal swabs (n = 7). In addition, two isolates of genotype B were to be found in throat of the lizards sampled. When looking at isolates of *M. granulomatis* genotype C and *M. viride*, the origin of isolation presented more divers. An overview of isolates, host species, origin of isolation and associated pathological findings is shown in Table 2.

**TABLE 2.**
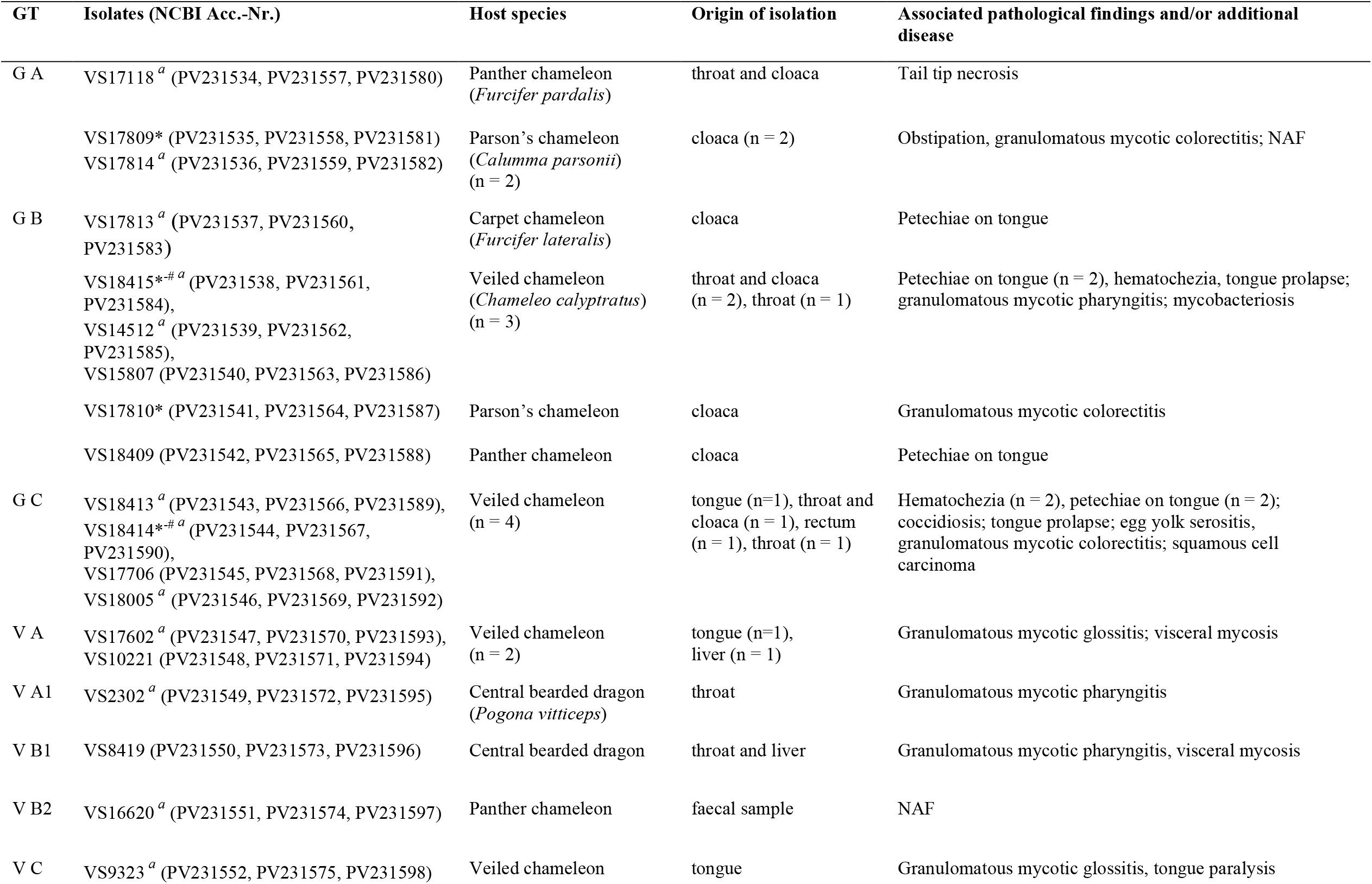
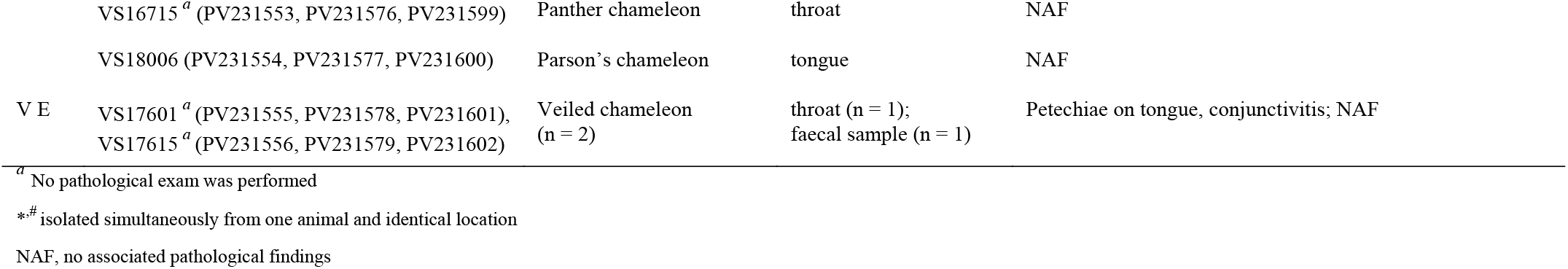
Summary of *Metarhizium granulomatis* and *M. viride* genotypes (GT) and isolates identified by sequence analysis of ribosomal DNA, host species, origin of isolate, associated pathological findings and/or additional disease.

| GT | Isolates (NCBI Acc.-Nr.) | Host species | Origin of isolation | Associated pathological findings and/or additional disease |
| --- | --- | --- | --- | --- |
| G A | VS17118 <sup>a</sup> (PV231534, PV231557, PV231580) | Panther chameleon<br>(*Furcifer pardalis*) | throat and cloaca | Tail tip necrosis |
|  | VS17809* (PV231535, PV231558, PV231581)<br>VS17814 <sup>a</sup> (PV231536, PV231559, PV231582) | Parson's chameleon<br>(*Calumma parsonii*)<br>(n = 2) | cloaca (n = 2) | Obstipation, granulomatous mycotic coloproctitis; NAF |
| G B | VS17813 <sup>a</sup> (PV231537, PV231560, PV231583) | Carpet chameleon<br>(*Furcifer lateralis*) | cloaca | Petechiae on tongue |
|  | VS18415*<sup>#a</sup> (PV231538, PV231561, PV231584),<br>VS14512 <sup>a</sup> (PV231539, PV231562, PV231585),<br>VS15807 (PV231540, PV231563, PV231586) | Veiled chameleon<br>(*Chameleo calyptratus*)<br>(n = 3) | throat and cloaca<br>(n = 2), throat (n = 1) | Petechiae on tongue (n = 2), hematochezia, tongue prolapse; granulomatous mycotic pharyngitis; mycobacteriosis |
|  | VS17810* (PV231541, PV231564, PV231587) | Parson's chameleon | cloaca | Granulomatous mycotic coloproctitis |
|  | VS18409 (PV231542, PV231565, PV231588) | Panther chameleon | cloaca | Petechiae on tongue |
| G C | VS18413 <sup>a</sup> (PV231543, PV231566, PV231589),<br>VS18414*<sup>#a</sup> (PV231544, PV231567, PV231590),<br>VS17706 (PV231545, PV231568, PV231591),<br>VS18005 <sup>a</sup> (PV231546, PV231569, PV231592) | Veiled chameleon<br>(n = 4) | tongue (n=1), throat and cloaca (n = 1), rectum (n = 1), throat (n = 1) | Hematochezia (n = 2), petechiae on tongue (n = 2); coccidiosis; tongue prolapse; egg yolk serositis, granulomatous mycotic coloproctitis; squamous cell carcinoma |
| V A | VS17602 <sup>a</sup> (PV231547, PV231570, PV231593),<br>VS10221 (PV231548, PV231571, PV231594) | Veiled chameleon<br>(n = 2) | tongue (n=1), liver (n = 1) | Granulomatous mycotic glossitis; visceral mycosis |
| V A1 | VS2302 <sup>a</sup> (PV231549, PV231572, PV231595) | Central bearded dragon<br>(*Pogona vitticeps*) | throat | Granulomatous mycotic pharyngitis |
| V B1 | VS8419 (PV231550, PV231573, PV231596) | Central bearded dragon | throat and liver | Granulomatous mycotic pharyngitis, visceral mycosis |
| V B2 | VS16620 <sup>a</sup> (PV231551, PV231574, PV231597) | Panther chameleon | faecal sample | NAF |
| V C | VS9323 <sup>a</sup> (PV231552, PV231575, PV231598) | Veiled chameleon | tongue | Granulomatous mycotic glossitis, tongue paralysis |
|  | VS16715<sup>a</sup> (PV231553, PV231576, PV231599) | Panther chameleon | throat | NAF |
|  | VS18006 (PV231554, PV231577, PV231600) | Parson's chameleon | tongue | NAF |
| V E | VS17601<sup>a</sup> (PV231555, PV231578, PV231601),<br>VS17615<sup>a</sup> (PV231556, PV231579, PV231602) | Veiled chameleon<br>(n = 2) | throat (n = 1);<br>faecal sample (n = 1) | Petechiae on tongue, conjunctivitis; NAF |

<sup>a</sup> No pathological exam was performed
<sup>\*#</sup> isolated simultaneously from one animal and identical location
NAF, no associated pathological findings

Pathological examination was performed on 7 of 23 (30.4%) of the lizards sampled. One isolate of *M. granulomatis* A as well as one of genotype B caused granulomatous colorectitis in Parson’s chameleons (n = 2). The same pathological findings were to be made in a veiled chameleon, caused by *M. granulomatis* genotype C. Three isolates of *M. granulomatis* genotype B were the cause of petechial haemorrhages on the tongue of three different chameleon species, furthermore genotype C caused the same finding in veiled chameleons (n = 2). *M. viride* genotype A and one isolate of genotype C were found as the cause of granulomatous glossitis in two veiled chameleons. Furthermore, one isolate belonging to *M. viride* genotype D and one belonging to genotype B were the cause of granulomatous mycotic pharyngitis in central bearded dragons (n = 2). Associated pathological findings were lacking in five of the animals presented (21.7%) (Table 2).

## 4. Discussion

*M. granulomatis* and *M. viride* are fungal pathogens, which are able to cause disease in various species of chameleons and bearded dragons (4). In the present study, fungal isolates were taken from six different locations of five different lizard species. This study presents a case of granulomatous colorectitis in a Parson’s chameleon, caused by *M. granulomatis*. This constitutes the first documented instance of this particular condition in this chameleon species. Furthermore, petechia on tongue, or a tail tip necrosis were seen in panther chameleons and a carpet chameleon infected with *M. granulomatis*. These findings suggest that *M. granulomatis* may have a broader host range and pathogenicity, as previously described (Schmidt et al., 2017b). Ongoing, *M. viride* was isolated from a healthy Parson’s chameleon for the first time. Even though there were associated pathological finding in 78.2% (18 of 23) of the lizards sampled, no significant correlations between reptile species, origin of isolation and pathological findings were to be made.

By sequencing of the SSU, ITS1-5.8S and LSU, Schmidt et al. (2017b) yielded five different genotypes of *M. granulomatis*, furthermore Schmidt et al. (2017a) assumed four different genotypes of *M. viride* by sequencing of the LSU. Sequencing of the RPB1 was not target-orientated to yield different genotypes of *M. granulomatis*, as there was no difference in amino acid structure. Also, the additional outgroup taxa clustered with *M. granulomatis* isolates, so that it appears that RPB1 is similar for *M. granulomatis* and the PARB-clade. Notwithstanding, a clear distinction between *M. viride* and *M. granulomatis* genotypes was possible by using this method. Four genotypes of *M. viride* could be distinguished regarding RPB1. The Maximum Composite Likelihood (MCL) approach showed that isolates of genotype A, D and the two isolates with previously unpublished genotype have the same amino acid structure and therefore, they cannot be differentiated by using only this dataset. *M. viride* genotype C isolates showed differences in amino acid structure and therefore clustered separately. Ongoing, there were significant differences that translated into amino acid structure of the two *M. viride* genotype B isolates, which leads to a renaming of the isolates into *M. viride* genotype B1 [NCBI Acc.-Nr. PV231550] and genotype B2 [NCBI Acc.-Nr. PV231551], with genotype B1 having the least percentage of similarity to other isolates (Suppl. 1).

Sequence analyses of RPB2 did not show any distinction between all of the fungal isolates. *M. granulomatis* as well as *M. viride* and also the PARB-clade clustered together, as there were no significant differences in amino acid structure. In conclusion, sequencing of the RPB2 alone is not practicable to analyze phylogenetic diversity (Suppl 2).

Similar results were obtained when analyzing the TEF dataset. The only isolates showing differences in amino acid structure and therefore clustering apart from others, were the ones belonging to *M. granulomatis* genotype A. All of the other isolates sequenced could not be differentiated using this method. However, isolates of the PARB-clade showed significant differences in amino acid structure of the TEF, compared to *M. granulomatis* and *M. viride* (Suppl. 3).

The multilocus dataset yielded three different genotypes of *M. granulomatis*. Genotype A, B and C showed differences in nucleotides that translated into three different amino acid structures. Diversity of the isolates was <98.7%. According to Stackebrandt & Ebers, 2006, who defined a threshold for species separation of ≥98.7–99%, in this case it would be appropriate to speak of subspecies. It was not possible to distinguish between *M. granulomatis* subspecies C and D, which leads to the conclusion that the isolates formerly belonging to genotype D (VS18413, VS18414) should be reclassified (Fig. 2).

Furthermore, six genotypes of *M. viride* could be yielded. By evaluating the multilocus dataset, it turns out that the isolate, which was formerly labelled as *M. viride* genotype D (VS2302), is genetically very similar to the isolates of genotype A, which clustered together. At this point, one should speak of a genetic divergence, based on the results of the Maximum Composite Likelihood (MCL) approach, which showed an identity of >98.7% to genotype A isolates. However, since the percentage of similarity to the other isolates of *M. viride* is <98.7%, it is suggested to reclassify the isolate VS2302. Being renamed as *M. viride* genotype A1, it appears to be a separate subspecies with genotype A.

The same applies to earlier mentioned isolates of *M. viride* genotype B1 (VS8419) and B2 (VS16620). The multilocus approach also underlies that they are genetically distinct and therefore supports a subdivision within genotype B. Further, the isolates belonging to *M. viride* genotype C clustered together and built a group of subspecies. The isolates of previously unpublished genotype (VS17601, VS17615) also clustered separately from others regarding amino acid structure. The proposal is to refer to them as *M. viride* genotype E. Finally, after sequencing and analyzing the data collected in this study, it can be postulated that an individual analysis of RPB1, RPB2 and TEF is not target-orientated for a specifying of *M. granulomatis* and *M. viride*. A multilocus analysis of the genes named above, however, represents a comprehensive and meaningful approach to differentiate between the species complexes *M. granulomatis* and *M. viride* and to furthermore define subspecies within these species’ complexes.

## Data Availability Statement

The NCBI Acc.-Nr. for the newly generated fragments of the RPB1 are PV231534, PV231535, PV231536, PV231537, PV231538, PV231539, PV231540, PV231541, PV231542, PV231543, PV231544, PV231545, PV231546, PV231547, PV231548, PV231549, PV231550, PV231551, PV231552, PV231553, PV231554, PV231555, PV231556.

For RPB2, the NCBI Acc.-Nr. are PV231557, PV231558, PV231559, PV231560, PV231561, PV231562,PV231563,PV231564,PV231565,PV231566,PV231567,PV231568, PV231569, PV231570, PV231571, PV231572, PV231573, PV231574, PV231575, PV231576, PV231577, PV231578, PV231579.

NCBI Acc.-Nr. for the TEF are PV231580, PV231581, PV231582, PV231583, PV231584, PV231585, PV231586, PV231587, PV231588, PV231589, PV231590, PV231591, PV231592, PV231593, PV231594, PV231595, PV231596, PV231597, PV231598, PV231599, PV231600, PV231601, PV231602.

